# Hormone circuit analysis explains why most HPA drugs fail for mood disorders and predicts the few that work

**DOI:** 10.1101/2024.06.10.598205

**Authors:** Tomer Milo, Shiraz Nir Halber, Moriya Raz, Avi Mayo, Uri Alon

**Affiliations:** Department of Molecular Cell Biology, Weizmann Institute of Science, Rehovot 76100, Israel

## Abstract

Elevated cortisol causes morbidity in chronic stress and mood disorders, including metabolic and cardiovascular diseases. There is therefore interest in developing drugs that lower cortisol by targeting its endocrine pathway, the hypothalamic-pituitary-adrenal (HPA) axis. Several promising HPA-modulating drugs have, however, failed to lower long-term cortisol in mood disorders such as major depressive disorder despite their effectiveness in situations where high cortisol is caused by a tumor (Cushing’s syndrome). Why these drugs failed is not well understood. Here we use a mathematical model of the HPA axis to show that the pituitary and the adrenal glands compensate for the effect of drugs by adjusting their functional mass, a feedback compensation that is absent in Cushing tumors. To find potential drug targets, we carried out a systematic *in silico* analysis of points of intervention in the HPA axis. We find that only two interventions that target corticotropin-releasing hormone (CRH) can lower long-term cortisol. Other drug targets either fail to lower cortisol due to gland-mass compensation or lower cortisol but harm other aspects of the HPA axis. Thus, we identify potential drug targets, including CRH-neutralizing antibodies and CRH-synthesis inhibitors, for lowering long-term cortisol in mood disorders and in those suffering from chronic stress. More generally, this study indicates that understanding the slow compensatory mechanisms in endocrine axes can be crucial in order to prioritize drug targets.

## Introduction

Cortisol is a steroid hormone produced by the adrenal glands in response to physical or psychological stressors. It acts on almost every tissue in the body and mediates the stress response by regulating metabolism, cognitive functions and immune responses.

Cortisol level is controlled by the hypothalamus-pituitary-adrenal (HPA) axis, a cascade of three hormones. In response to stressor inputs, the hypothalamus secretes corticotropin-releasing hormone (CRH). CRH stimulates the secretion of adrenocorticotropic hormone (ACTH) by corticotroph cells in the anterior pituitary. ACTH in turn signals the adrenal cortex to secrete cortisol. Cortisol inhibits the production and secretion of the two upstream hormones, CRH and ACTH ^1^, forming a negative feedback loop.

Prolonged elevated levels of cortisol, a condition known as hypercortisolism, can lead to a range of health issues ^1^. These include weight gain, high blood pressure, diabetes, osteoporosis, muscle weakness, thinning skin, increased bruising, slower wound healing, and mood changes. Additionally, hypercortisolism can cause disruptions in sleep and memory, reduce libido, and compromise the immune system, making the body more susceptible to infections ^2–4^.

Hypercortisolism occurs in the context of chronic stress, such as that associated with low socioeconomic status ^5,6^ and also in mood disorders such as major depressive disorder (MDD) ^7^ and bipolar disorder (BD) ^8,9^. Hypercortisolism can also be caused by drugs and tumors. Drugs such as glucocorticoid steroids are cortisol analogues that cause the above mentioned health issues upon prolonged treatment. Tumors in Cushing syndrome escape HPA regulation and cause elevated cortisol. Cushing’s syndrome is often treated by tumor removal. When surgery is not possible in Cushing patients, the negative health effects of high cortisol are treated by cortisol-modulating drugs such as cortisol synthesis inhibitors and cortisol receptor antagonists ^10,11^.

Interestingly, Cushing’s tumors and corticosteroids frequently cause mood episodes ^12–16^. This causal effect of elevated chronic cortisol on mood, as well as the strong association of stress and MDD, has raised the hope that cortisol-lowering drugs could improve mood disorder symptoms in MDD and BD ^17–19^

It thus came as a disappointment when HPA medications useful in Cushing’s syndrome failed in clinical trials for MDD and BD (see Table 1) ^20^. These drugs include cortisol- and ACTH-synthesis inhibitors and glucocorticoid receptor (GR) antagonists. Drugs that lowered cortisol in Cushing syndrome failed to lower long-term cortisol in people with mood disorders, and some drugs such as GR antagonists even raised cortisol levels. Why HPA drugs that lower cortisol in Cushing syndrome show limited efficacy in mood disorders is not well understood. It is of interest to explore whether still untested HPA drugs might lower cortisol in the context of chronic stress and mood disorders.

**Table 1.**
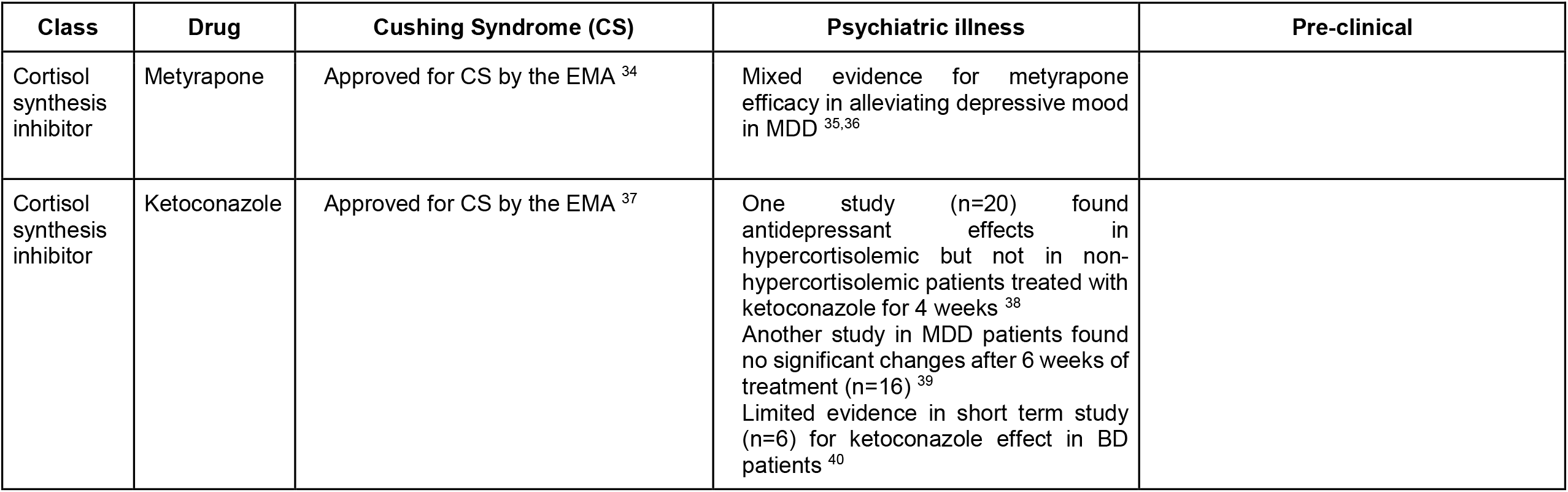
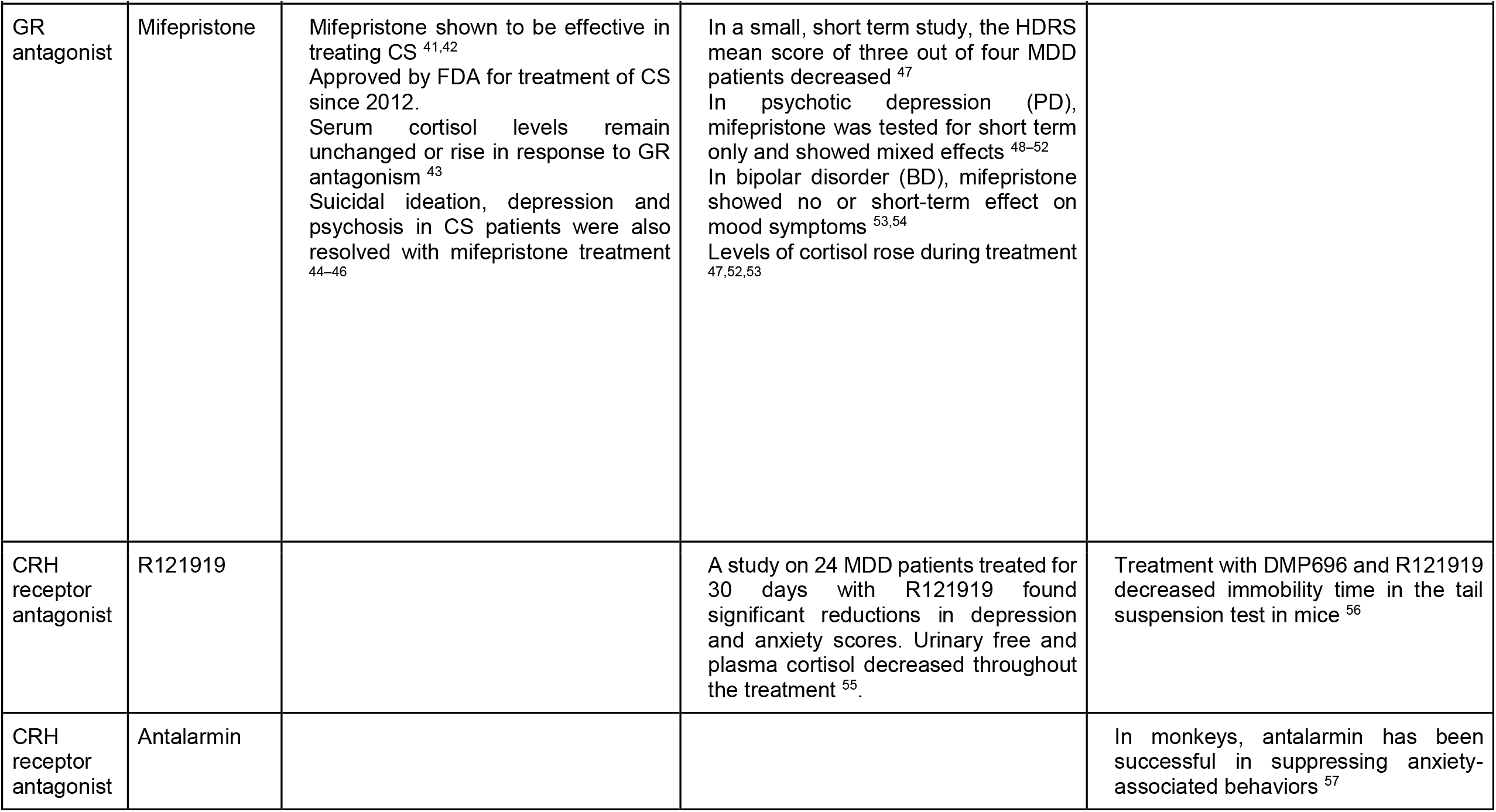
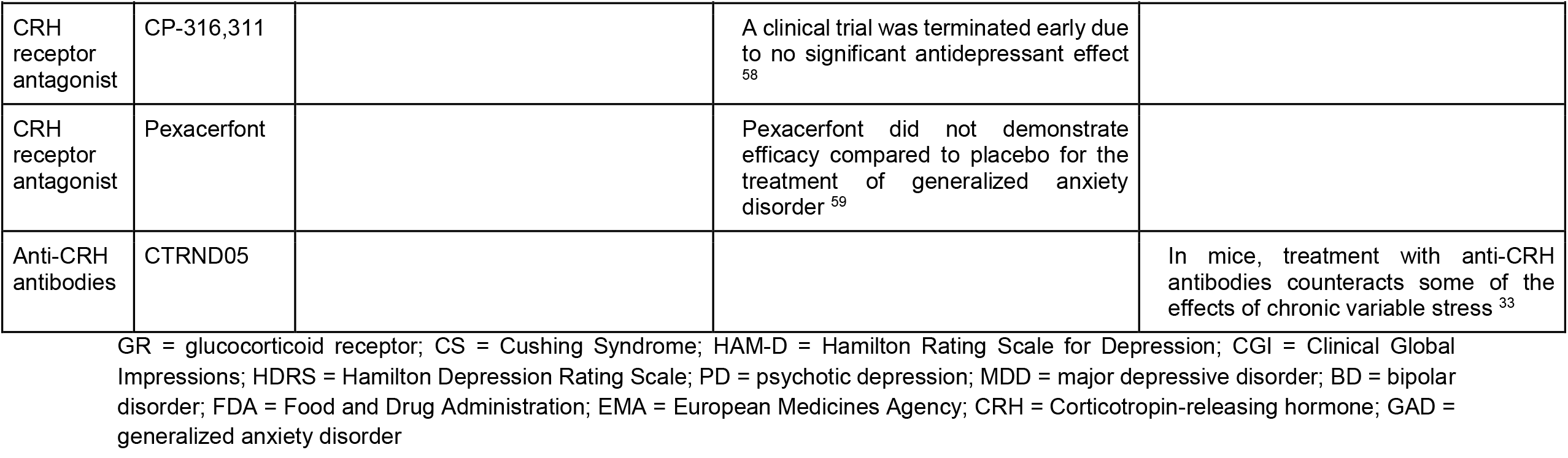
Long-term efficacy of HPA-related drugs in hypercortisolism conditions.

**Table 2.** Definitions of the HPA mathematical model variables and parameters.

| Dynamic variable | Definition |
| --- | --- |
| $x_1$ | Corticotropin-releasing hormone (CRH) |
| $x_2$ | Adrenocorticotrophic hormone (ACTH) |
| $x_3$ | Cortisol hormone |
| $P$ | Pituitary corticotroph functional mass |
| $A$ | Adrenal functional mass |
| Parameter | Definition |
| $b_1, b_2, b_3$ | CRH, ACTH, cortisol production rates |
| $a_1, a_2, a_3$ | CRH, ACTH, cortisol degradation rates |
| $b_P, b_A$ | Growth rates of pituitary corticotroph and adrenal functional mass |
| $a_P, b_P$ | Removal rates of pituitary corticotroph and adrenal functional mass |
| $K_{GR}$ | Glucocorticoid receptor dissociation constant |
| Intervention | Definition |
| $u$ | Stressor input |
| $I_1, I_2, I_3$ | CRH, ACTH and cortisol synthesis inhibitors |
| $C_1, C_2, C_3$ | CRH, ACTH and cortisol receptor antagonists |
| $A_1, A_2, A_3$ | CRH, ACTH and cortisol neutralizing antibodies |

Here we address this using a systems pharmacology approach, by employing a recent advance in mathematical modeling of the HPA axis. The new model ^21^ updated the classical HPA model which works on the timescale of the hormone lifetimes, namely minutes to hours ^22^. The classical model is thus not suited to address the timescale of weeks to months needed to assess chronic cortisol levels. The weeks-months timescale was introduced in the Karin et al mathematical model of the HPA axis by including changes over time in the functional mass of the endocrine glands ^21^. The larger the gland mass, the more hormone it secretes per unit input hormone. The gland size in the model is governed by well-characterized interactions which had not been previously considered on the system level, namely that gland mass is regulated by the HPA hormones that act as growth factors. CRH acts as a trophic factor for the corticotrophs of the pituitary ^23,24^ and ACTH serves as a growth factor for the cortisol-secreting cells in the adrenal ^25,26^. Since the cell turnover time in the pituitary and adrenal glands is on the order of months, glands grow and shrink on this slow timescale. The model shows how the gland masses adjust over months to buffer variation in physiological parameters, a property called *dynamical compensation* ^27^.

The gland-mass model was tested and validated using longitudinal hair cortisol measurements in healthy individuals ^28^ and in people with bipolar disorder ^9^ where it explained year-scale cortisol fluctuations. It was also validated and calibrated on a wide range of long-term phenomena such as hormone seasonality ^29^, recovery from chronic stress ^21^ and addiction ^30^. The model was extended to understand the timescales of MDD ^31^ and BD ^9^. The concept of changeable gland mass was also adapted to the thyroid axis to explain dynamics of thyroid diseases ^32^. These studies motivated us to use the HPA gland mass model to understand which HPA-modulating drugs might lower cortisol and which are destined to fail.

Here we use the gland-mass mathematical model to systematically test *in silico* many possible HPA interventions for lowering long-term cortisol. We find that most drugs do not lower cortisol in an intact HPA axis due to the compensatory capacity of the gland masses, which change over weeks to completely nullify the drug effect. This compensation is broken in Cushing tumors which escape HPA regulation, explaining why Cushing’s drugs are effective in lowering cortisol. We identify two CRH-associated drug targets that are expected to lower long-term cortisol in chronic stress conditions and mood disorders but not in Cushing’s syndrome. These drugs also preserve all HPA hormone levels and response features. This study thus proposes that certain CRH-modulating drugs, such as neutralizing anti-CRH antibodies ^33^, may be effective to lower long-term cortisol in stress-related disorders. More generally, this study indicates that understanding the slow compensatory mechanisms in endocrine axes can be crucial in order to prioritize drug targets.

## Results

### Under chronic stress, CRH-associated interventions normalize long-term cortisol, whereas other interventions are compensated and fail

To search for strategies to lower long-term cortisol levels, we used the HPA gland-mass model ^21^. The model incorporates the classical hormone cascade and negative feedback loop ^60^. The gland-mass interactions added to the classical HPA model are highlighted in bold in Figure 1A.

**Figure 1.**
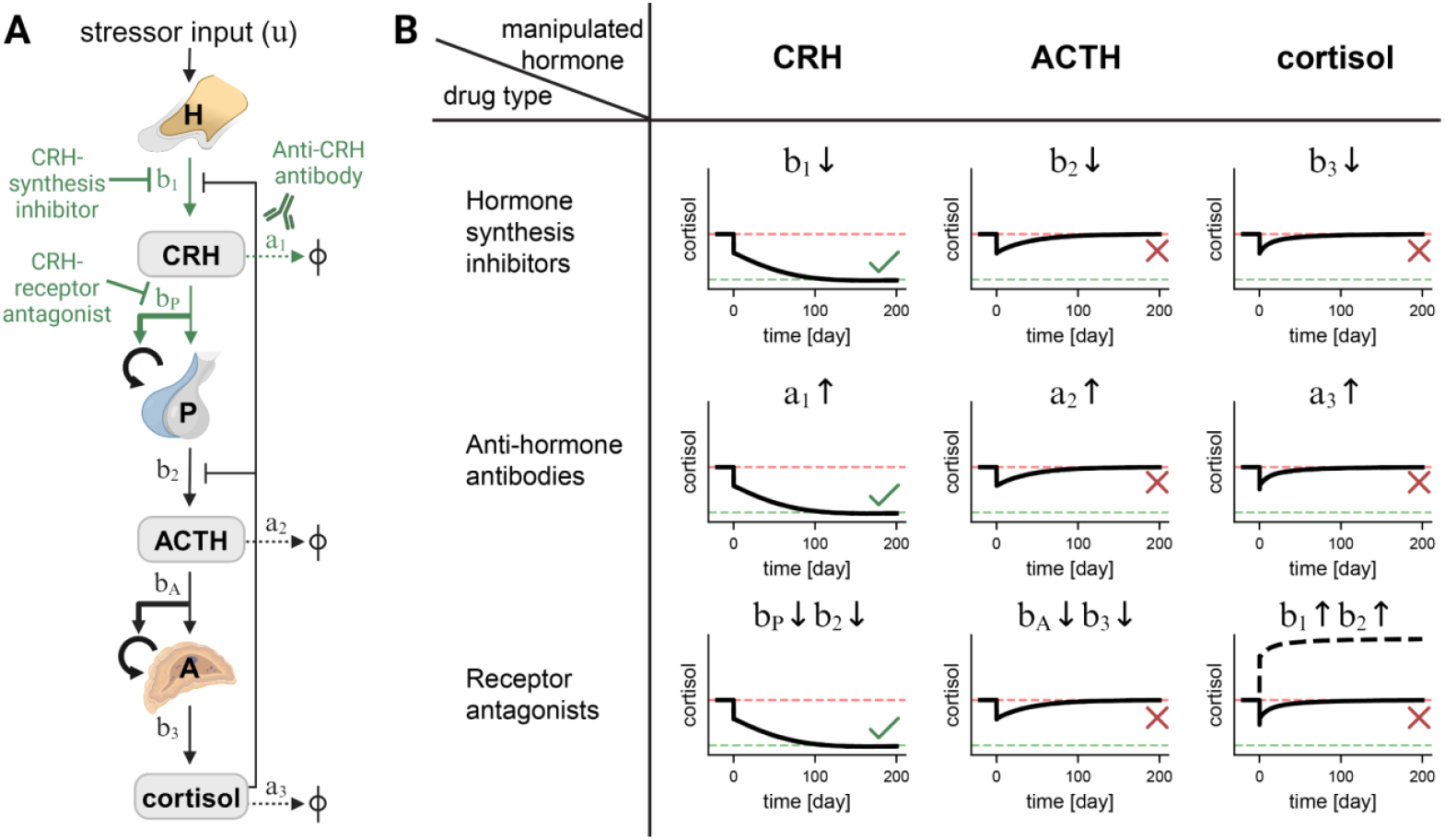
A circuit-to-target approach to lowering long-term cortisol levels in chronic stress conditions points to a few drug targets that can be effective, wheras most are predicted to fail. (**A**) The HPA circuit diagram. The hypothalamus H secretes CRH at rate *b*_1_ in response to a stressor input *u*. CRH causes the pituitary P to secrete ACTH at rate *b*_2_ and to grow in effective mass at rate *b*_*P*_. ACTH signals the adrenal gland to secrete cortisol at rate *b*_3_ and to grow in effective mass at rate *b*_*A*_ rate. The hormone removal rates are *a*_1_, *a*_2_ and *a*_3_ for CRH, ACTH and cortisol, respectively. Thick arrows indicate the growth factor interactions added in the Karin et al model that affect gland sizes on the scale of months. The drugs predicted to be effective and their points of intervention are illustrated in green. (**B**) Simulations of HPA interventions. Simulations began with an elevated level of cortisol (marked by horizontal dashed red lines) due to chronic stress input and were run for 200 days. A single simulated drug was administered at time zero. The parameter influenced by each drug and its direction of change is indicated in each panel (see Methods). As an example, note that cortisol receptor antagonists inhibit the negative feedback of cortisol on CRH and ACTH synthesis and thus effectively increase CRH and ACTH production rates, *b*_1_ and *b*_2_, respectively. Simulations that succeeded in reducing cortisol to a normal level (horizontal dashed green lines) are marked with a green check mark, whereas those that failed are marked with a red X. In the case of cortisol receptor antagonists, the dashed black line indicates cortisol level, and the continuous black line is the cortisol net effect through GR signaling on target cells after accounting for the receptor blocking effect by the antagonists.

In these interactions, the dynamics of the pituitary and adrenal masses are under the control of their upstream HPA hormone growth factors, CRH and ACTH, respectively. Since the model was found to be accurate in a wide range of clinical situations that undergo changes over months ^9,21,28–31^, we reasoned that it would be informative also for drug effects.

We tested *in silico* all possible points of intervention and asked whether they reduce cortisol in the long term. In this “circuit-to-target” approach, we systematically modeled each intervention’s effect on the model parameters and looked for those that reduced steady-state cortisol (see Methods). In this way, we simulated agonists and antagonists for cortisol, ACTH or CRH receptors, antibodies against each of the three hormones, and synthesis inhibitors of each of the three hormones (Figure 1B).

We also tested possible combinations of interventions. We found that no new effective combinations arise apart from combinations of the single drugs described next.

We find that after a transient period of a few weeks, the HPA glands in the model change in size to compensate for most of the possible interventions. For example, blocking the glucocorticoid receptor (GR) with GR antagonists led to an increase in the adrenal cortex mass. The increased adrenal mass generated higher levels of cortisol that *precisely negated* the reduction in GR binding to cortisol (Figure 1B, bottom right panel). Thus, a GR antagonist has no net effect on GR signaling after a transient period of a few weeks. The rise in cortisol agrees with observations from clinical trials using the GR antagonists (Table 1) ^47,52,53^. This effect would not be seen in the classical HPA model.

Similarly, ACTH receptor antagonists did not affect cortisol levels after a few weeks because the adrenal gland grew to compensate precisely for the inhibition. Synthesis inhibitors of cortisol and ACTH likewise had only a transient effect of a few weeks, which vanished once gland masses changed to fully compensate for the intervention.

It can be shown mathematically why the compensatory properties of the circuit prevent most parameter changes from altering cortisol steady-state in the long term (see equations (6)-(15) in Methods). This is because cortisol steady state, *cortisol*_*st*_, is robust to changes in most of the circuit parameters. It depends on only a few parameters that associate with CRH signaling:

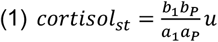

Crucially, this equation also reveals which interventions can lower cortisol steady state. According to equation (1), cortisol steady state can be reduced by CRH-related interventions (Figure 1B, left column). First, one can lower CRH production rate *b*_1_ by using CRH-synthesis inhibitors. Second, one can lower the effect of CRH on pituitary corticotroph cell growth rate *b*_*P*_. This can be done by inhibiting the CRH-receptor on the pituitary corticotrophs using a receptor antagonist ^55^. Another way to lower cortisol is to increase the CRH removal rate *a*_1_, for example by using antibodies that bind and neutralize CRH. An anti-CRH antibody has indeed been shown to suppress the HPA axis in stressed mice ^33^. Such drugs may be candidates for treating chronic-stress conditions.

The model further predicts that lowering the input *u*, the stress signal in the brain communicated to the hypothalamus and leading to CRH secretion, can also lower cortisol steady state. This might relate to psychotherapy, exercise and other lifestyle interventions that reduce stress ^61^. Finally, increasing the pituitary corticotroph removal rate *a*_*P*_ should also lower cortisol steady state.

All other drugs that target ACTH or cortisol, are predicted to have only a transient effect lasting a few weeks. This includes receptor antagonists, hormone production inhibitors, or anti-hormone antibodies. Such drugs do not affect cortisol steady state and thus fail to lower cortisol levels in the long term (Figure 1B, middle and right columns). The classical HPA model with nonadjustable glands predicts that cortisol steady state would depend on ACTH and cortisol parameters (see Figure S1), and thus cannot predict these compensation effects.

Importantly, these conclusions on drug effect do not depend on the HPA model parameter values, since they can be analytically derived from the model’s steady state solution. They are thus a robust prediction.

### CRH-synthesis inhibitors and anti-CRH antibodies normalize all HPA hormones and glands

Next, we asked whether the effective drugs normalize not only cortisol but also the levels of the other HPA components. Two of the CRH interventions, namely CRH-synthesis inhibitors and anti-CRH antibodies, are predicted to normalize the entire HPA axis - all hormone levels and gland masses return to normal (Figure 2 A-B).

**Figure 2.**
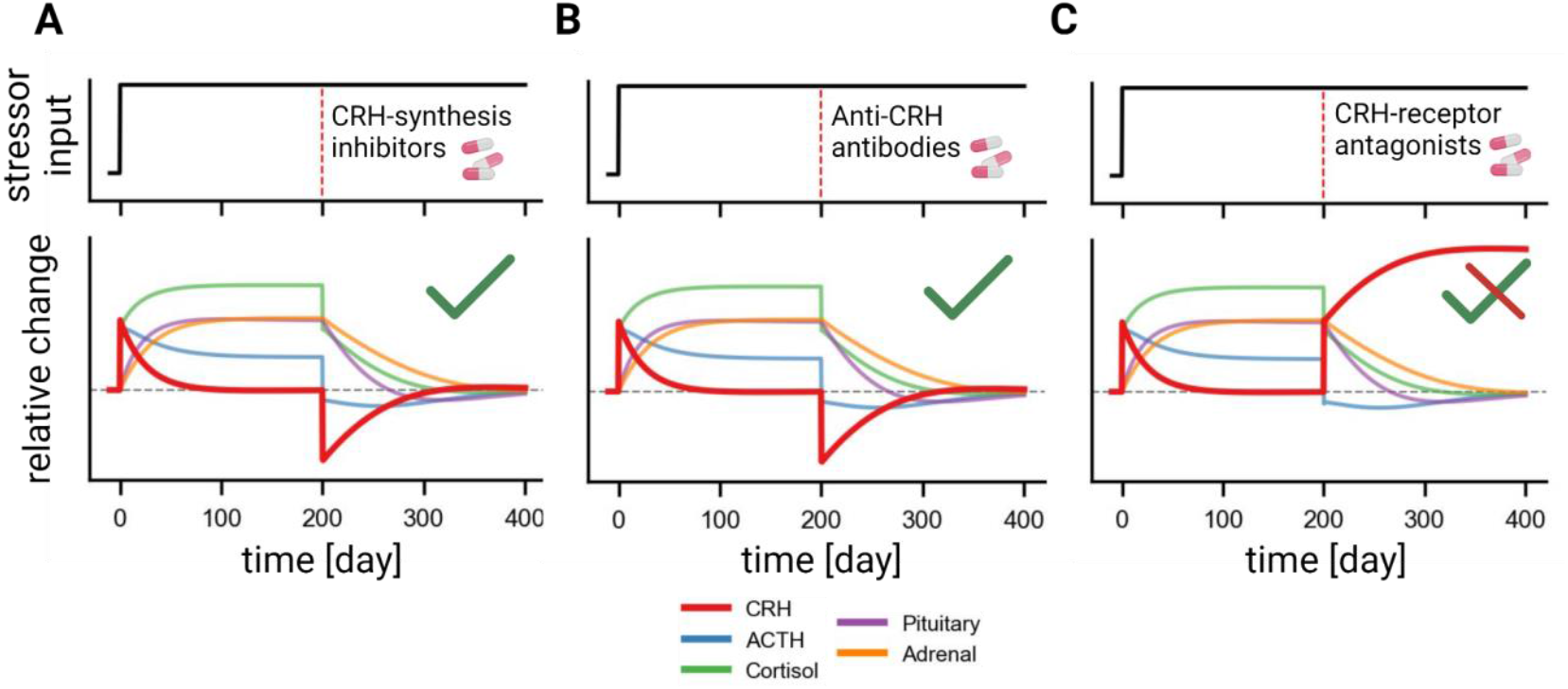
Two of the CRH-targeting drugs normalize all hormone and gland masses in chronic stress, whereas one raises CRH. Chronic stress was simulated by increasing the input *u* at time zero and keeping it elevated for the entire simulation (upper panels). The drug was administered 200 days later (red vertical dashed line). The simulated HPA responses to CRH-synthesis inhibitors (**A**), anti-CRH antibodies (**B**) and CRH-receptor antagonists (**C**) are presented in the lower panels. CRH dynamics are highlighted in red. The gray horizontal dashed line indicates the healthy baseline.

In contrast, CRH-receptor antagonists are predicted to reduce long-term cortisol but increase long-term CRH levels (Figure 2C). Higher CRH steady state is due to the pituitary’s integral feedback loop that locks CRH levels ^21,62^. Physiologically, CRH-receptor antagonists reduce the growth-factor effect of CRH on pituitary cells, and thus more CRH is needed to keep the pituitary at a fixed size.

### CRH interventions preserve acute responses to relative stressors

Given the predicted efficacy of CRH-synthesis inhibitors and anti-CRH antibodies in normalizing the HPA axis under chronic stress conditions, we next asked whether these interventions also preserve acute stress responses. It is important to evaluate the acute stress response, in order to avoid drugs that normalize the axis in the long term, but impair short-term responses on the timescale of hours which are critical to successful stress responses.

We simulated the HPA model with acute stressors in the form of short-term pulses of input *u* that are 2 times higher than the baseline input (Figure 3, top row). The first acute stressor input was simulated at the healthy baseline and serves as a reference for comparison. The second acute stress pulse was simulated during the chronic stress period with drug treatment (Figure 3, top row).

**Figure 3.**
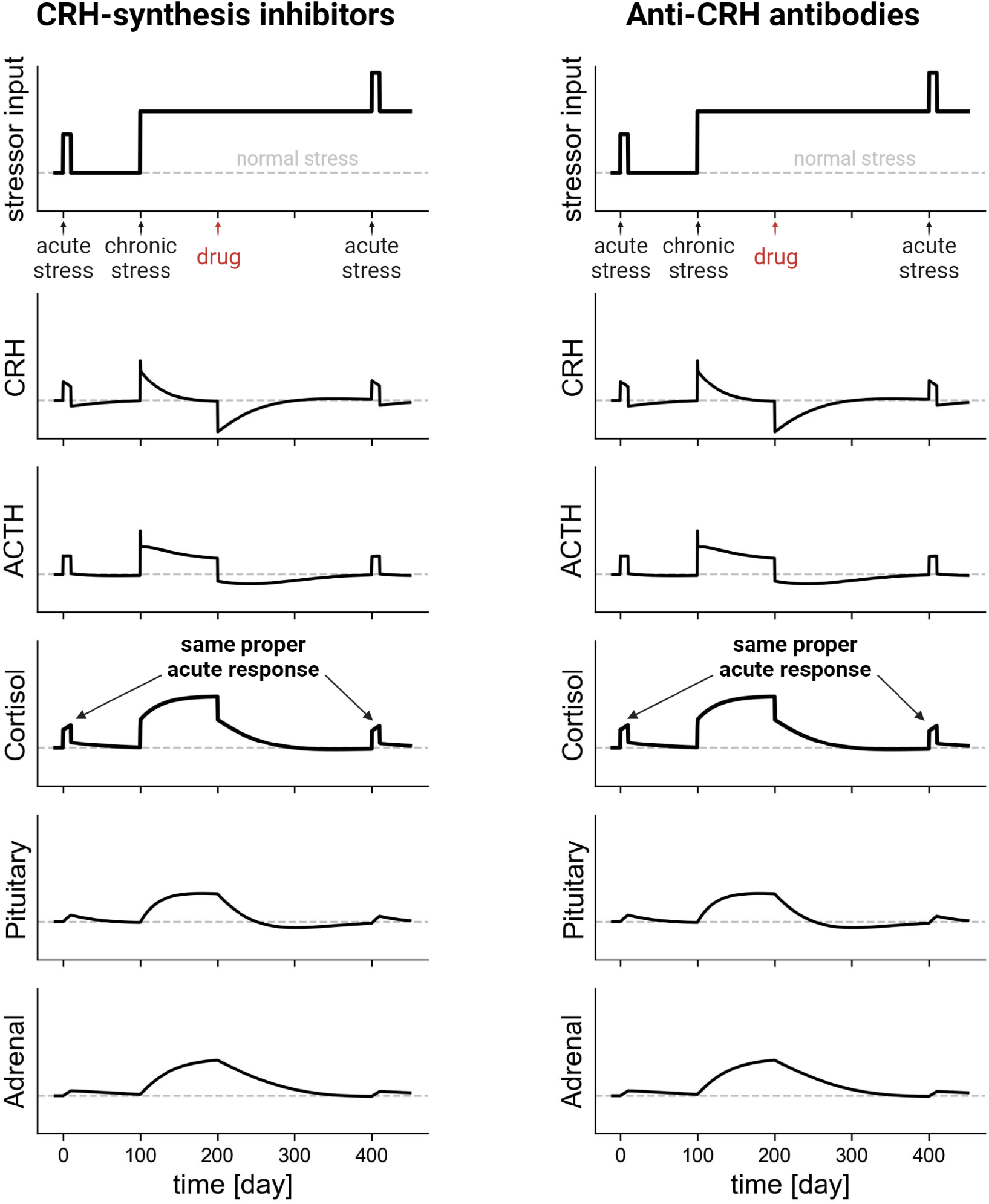
HPA responses to acute stress are preserved during CRH-related interventions. Acute stress was simulated as a 2-fold, short-term pulse above a baseline (top row). The first acute stress was introduced during the healthy period, before the chronic stress. The second acute stress was simulated during the chronic stress period, several months after drug treatment began. The response of HPA hormones and glands to these stressors for CRH-synthesis inhibitors (left column) or for anti-CRH antibodies (right column) is depicted in the bottom five rows. The dashed lines indicate the healthy normal state.

We find that the HPA acute stress response under the CRH-targeting medications were identical to the normal healthy responses (Figure 3). In line with this prediction, a proper acute stress response was observed in mice treated with anti-CRH antibodies after two weeks of chronic variable stress ^33^. We thus conclude that CRH-synthesis inhibitors and anti-CRH antibodies are both predicted to normalize HPA components in the long term and preserve relative acute stress responses.

### Endogenous Cushing’s syndrome responds to different drug targets due to loss of gland compensation

We next aimed to understand why HPA-targeting drugs that failed in MDD and BD trials succeed in treating hypercortisolism in Cushing’s syndrome. We investigated the two main classes of Cushing’s syndrome - tumors that produce an excessive amount of ACTH (Cushing’s disease and ectopic ACTH syndrome) and tumors that overproduce cortisol (adrenal gland tumors). The ACTH-dependent cases are more common ^63^. Importantly, the tumors escape the HPA feedback loops - high cortisol does not suppress hormone production in the mutant tumor cells.

Cushing’s tumors are often treated with surgery. However, tumor location, patient health status, or patient preference can favor medication rather than surgery. Drugs are also sometimes used temporarily to control cortisol deleterious effects before surgery or while waiting for the effects of radiation therapy ^10,11^.

To analyze HPA drugs in Cushing’s syndrome, we follow the analysis of ^64^. We modeled the tumor secretion rates by adding the appropriate production terms to the equations (Methods, equations (17) and (25)). We modeled tumor growth by a logistic function ^65,66^ (Figure 4A-B, top panels). Our findings are not sensitive to the exact functional form of tumor growth. In both adrenal and pituitary adenomas, as long as the tumor is below a certain threshold of secretion rate (vertical dashed gray lines, Figure 4A-B), the HPA glands compensate to keep the hormones at normal levels (see Methods) ^64^. During this stage, the tumor is predicted to be subclinical with no overt symptoms.

**Figure 4.**
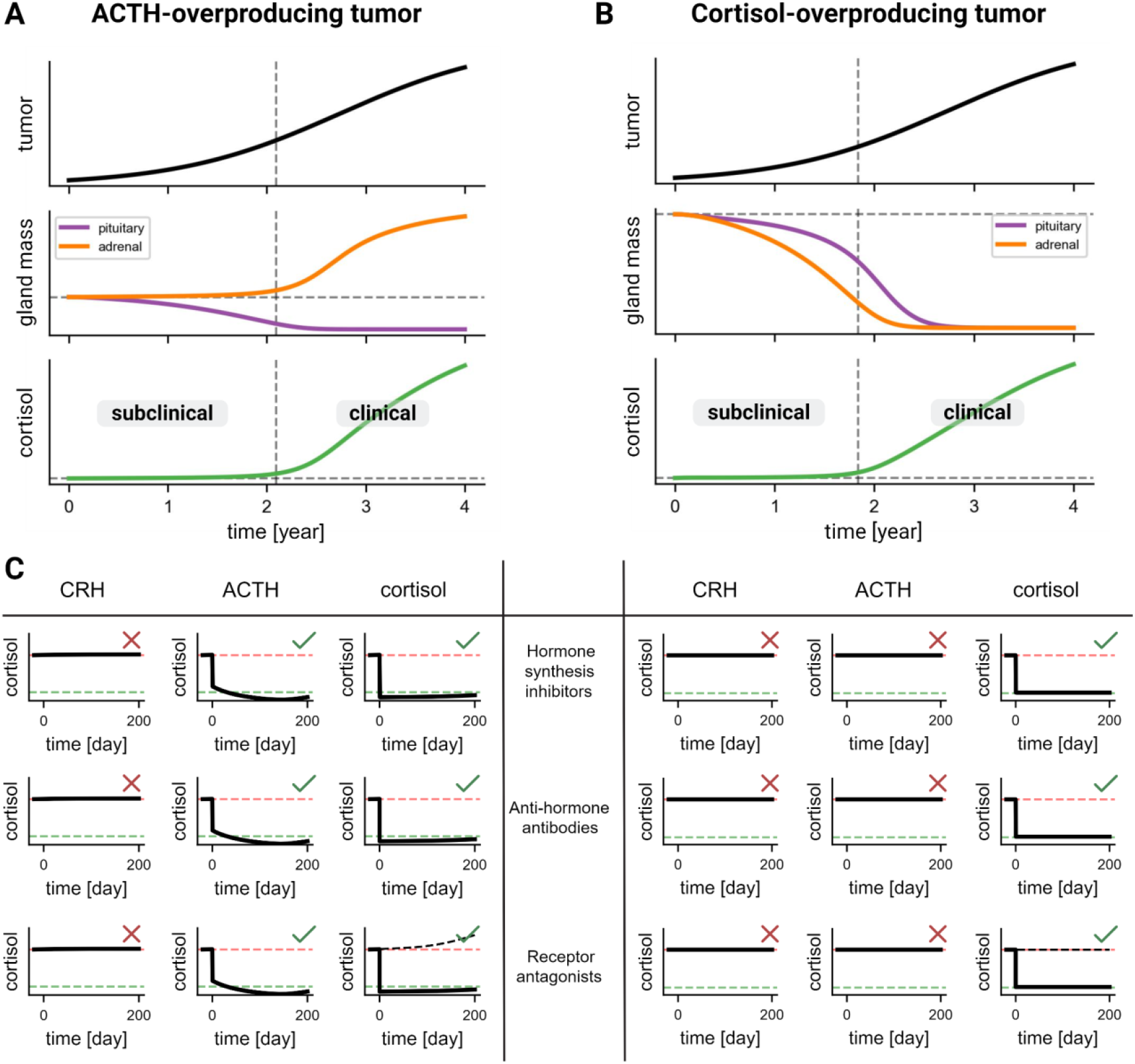
Dynamics of natural and medicated Cushing’s syndrome show that different sets of drugs are effective. (**A**-**B**) Simulations of ACTH-overproducing pituitary adenoma (**A**) and cortisol-overproducing adrenal adenoma (**B**). The HPA glands change their mass (middle panel) as the tumor grows (upper panel) and thus maintain homeostatic levels of cortisol (bottom panel) up to a threshold tumor secretion rate (vertical gray dashed line). Below this threshold cortisol levels do not rise - the subclinical phase. Above this threshold the glands cannot compensate and cortisol levels rise - the clinical phase. (**C**) Simulations of potential HPA-modulating drugs to treat pituitary adenoma (left table) and adrenal adenoma (right table). Red X marks a drug predicted to fail in treating Cushing, while a green check mark marks drugs predicted to work. Red and green horizontal dashed lines indicate abnormal and normal cortisol levels. In simulations of cortisol-receptor antagonists, we show cortisol levels in black dashed line and the net effect of cortisol on target cells, after the antagonist effect, in black continuous line.

In an ACTH-overproducing adenoma, the healthy pituitary corticotroph mass shrinks (Figure 4A, middle panel). Once the tumor secretion crosses a threshold, the pituitary is too small to compensate, and ACTH rises. An increase in ACTH leads to an increase in cortisol and to adrenal growth. In this regime, cortisol stead-state loses its normal independence on ACTH and cortisol parameters (see Methods). Thus, ACTH- or cortisol-targeting drugs can reduce cortisol levels (Figure 4C, left table). These include ACTH or cortisol synthesis inhibitors, ACTH or cortisol receptor antagonists, and anti-ACTH- or anti-cortisol-blocking antibodies.

The model also predicts that if the pituitary corticotrophs were not completely suppressed at the time of drug administration, the entire HPA axis could potentially recover apart from normalizing cortisol. This recovery happens because the drugs effectively increase the subclinical-clinical threshold and move the patient back into the subclinical regime (see Methods). Although all ACTH- and cortisol-targeting drugs lower long-term cortisol (Figure 4C, left table), only ACTH-synthesis inhibitor and anti-ACTH-blocking antibodies are predicted to normalize all HPA components (Supplemental Figure 2).

In an adrenal tumor, the healthy (non-tumor) adrenal glands shrink to compensate for the tumor’s overproduction of cortisol ^67,68^ (Figure 4B, middle panel). Once the healthy adrenal tissue is too small to compensate, cortisol level rises and inhibits the upstream hormones. As a result, the pituitary shrinks as well. At this stage, cortisol level is determined by the adrenal tumor alone (see Methods). Therefore, only drugs that manipulate cortisol parameters - cortisol synthesis inhibitors, anti-cortisol antibodies or cortisol receptor antagonists - should work to lower long-term cortisol levels (Figure 4C, right table). Similar to the ACTH-overproducing tumor, these cortisol-targeting drugs are predicted to increase the subclinical-clinical threshold and recover all HPA components (Supplemental Figure 2). Interestingly, Mifepristone, a GR antagonist, resolved psychosis and depression symptoms in Cushing’s patients ^44,45^ but was less efficient in MDD patients (Table 1), aligning with our results.

Note that in Cushing scenarios, drug effects are predicted to work on the timescales of hours since the effects do not depend on the slow gland timescales. Notably, different sets of drugs work for the two types of Cushing syndrome, and none of these are predicted to lower cortisol in the HPA axis without a tumor.

## Discussion

We present a systems pharmacology approach to discover HPA drug targets to lower long-term cortisol in mood disorders and chronic stress conditions. We used a validated mathematical model that takes into account compensatory changes in endocrine gland functional mass ^21^. Most drugs are predicted to fail due to compensation by gland changes over weeks. We identify two drug targets that are predicted to lower cortisol - CRH-synthesis inhibitors and anti-CRH antibodies. These interventions also normalize all other HPA hormones and maintain a proper response to stimuli. Other interventions that target ACTH or cortisol directly are predicted to transiently lower cortisol for several weeks, followed by a return of cortisol to its aberrant baseline. We also evaluated drug targets in Cushing’s tumors which bypass the normal HPA feedback loops. Thus, we conclude that CRH synthesis inhibitors or neutralizing antibodies cannot be compensated by the HPA axis, and are candidates to lower cortisol in mood disorders and in chronic stress.

These findings explain the failure of clinical trials that attempted to use HPA-modulating drugs such as GR antagonists and cortisol-synthesis inhibitors to treat mood disorders (Table 1). These same drugs, such as the GR antagonist mifepristone, work well in improving symptoms in Cushing’s syndrome patients ^10^. We propose that the failure of Cushing drugs in mood disorders can be understood by considering the ability of the HPA glands to compensate for these interventions by changing their functional mass. For example, administering GR antagonists reduces cortisol negative feedback on the upstream hormones, CRH and ACTH. Therefore, their levels rise and lead to higher secretion of cortisol and growth of the glands to exactly negate the GR antagonist effect on the receptor. In Cushing’s syndrome, due to tumors that have lost the HPA feedback loops, ACTH- and cortisol-targeting drugs work because the glands lose their ability to compensate for the drug effects. Note that the classical model, with no changes in gland masses, cannot explain the compensation of drugs by the HPA axis.

To systematically test all possible points of intervention in the HPA axis we extended the Karin et al mathematical model for the HPA axis which incorporates the slow timescale dynamics of the endocrine glands ^21,64^. We used numerical simulations to analyze HPA dynamics under different medication regimes. We also analytically solved the steady states of the system. This approach rigorously defines the potential points of intervention. The qualitative conclusions on which drugs work does not depend on the model parameter values.

Our model is a simplified representation of the complex HPA biology. Future work might consider modeling the implications of exact drug pharmacokinetics and a more accurate model of tumor growth. A dynamical aspect that we neglected but could be important to investigate is the diurnal pulsatile secretion pattern of CRH and ACTH ^69,70^. A more complete but more complex model can also include crosstalks with other endocrine systems such as the thyroid axis ^71,72^. Future work can consider treatment for other HPA related conditions such as congenital adrenal hyperplasia, Addison’s disease and post-traumatic stress disorder (PTSD).

In conclusion, our circuit-to-target approach explains why attempts to lower long-term cortisol using most HPA-targeting drugs are doomed to fail due to compensation by functional gland mass changes. We find that only a few drug targets can lower cortisol in the long term while preserving all other hormone levels. Predicted effective drugs include inhibitors of CRH synthesis or drugs that increase CRH removal such as anti-CRH antibodies. Such drugs may hold promise to treat cortisol-related mood disorders ^7–9,31^ and to mitigate the deleterious health effects of cortisol in those suffering from chronic stress. More generally, this study indicates that understanding the slow compensatory mechanisms in endocrine axes can be crucial in order to prioritize drug targets.

## Methods

### The HPA gland mass model

To test the effect of HPA-targeting drugs in chronic stress conditions and Cushing tumors, we used the HPA gland mass model ^21^ and added explicit parameters for each possible intervention:

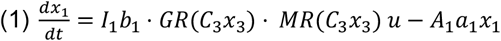

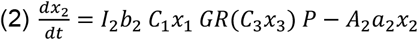

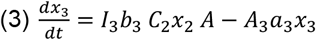

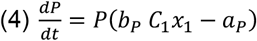

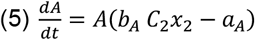

In response to an input stressor, *u*, the hypothalamus secretes CRH, *x*_1_, at a rate *b*_1_. CRH stimulates the corticotrophs at the pituitary, *P*, to secrete ACTH, *x*_2_, at a rate *b*_2_. ACTH signals the adrenal cortex of the two adrenal glands, whose total functional mass is *A*, to secrete cortisol, *x*_3_, at a rate *b*_3_. Cortisol inhibits the production of CRH and ACTH through the mineralocorticoid and glucocorticoid receptors, given by 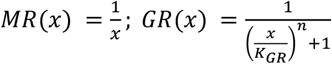, respectively. CRH, ACTH and cortisol degrade at rates *a*_1_, *a*_2_ and *a*_3_ respectively. The gland-mass model includes the trophic effects of CRH on the pituitary (*b*_*P*_*x*_1_) and of ACTH on the adrenal (*b*_*A*_*x*_2_).

We added an explicit parameter for each possible intervention: hormone-synthesis inhibitors, *I*_*i*_ < 1, reduce hormone production rate, *b*_*i*_, and does not affect anything else explicitly because *I*_*i*_ appears only multiplying *b*_*i*_; anti-hormone-blocking antibodies, *A*_*i*_ > 1, increase hormone removal rate, *a*_*i*_; hormone-receptor antagonists or agonists, *C*_*i*_, modulate the effect of hormone *x*_*i*_ on its corresponding receptor and thus are coupled to obtain the hormone net effect, *C*_*i*_*x*_*i*_. CRH-receptor antagonists or agonists, *C*_1_, affect both ACTH production rate, *b*_2_, and corticotroph growth rate, *b*_*P*_. Similarly, ACTH-receptor antagonists or agonists, *C*_2_, effectively modulate cortisol production rate, *b*_3_, and adrenal growth rate, *b*_*A*_. Note that agonists or antagonists to cortisol receptors affect CRH and ACTH production rates, *b*_1_ and *b*_2_, because cortisol inhibits CRH and ACTH synthesis in the hypothalamus and the pituitary, correspondingly.

### Analytical solutions of the HPA model’s steady state

#### Acute and chronic stress conditions

Physiological and psychological stressor inputs to the hypothalamus cause the secretion of CRH. We denote stressor input magnitude to the hypothalamus by *u*. Acute and chronic stressors are modeled as a short or a prolonged fold-change increase from the baseline input, *u* = 1. Input stressors that are very large induce high levels of cortisol that saturate the GRs, therefore we assume *x*_3_ ≫ *K*_*GR*_. Under this approximation and the GR Hill coefficient, *n* = 3, 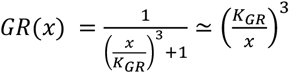. We also validated our conclusions with numerical simulations, relieving this assumption. In this regime the model and its steady state, denoted with *st* subscript, are:

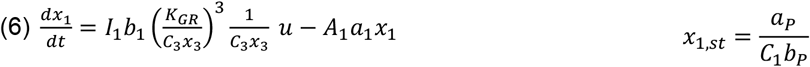

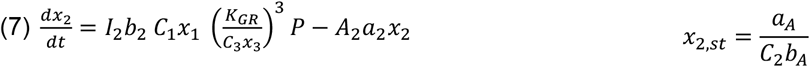

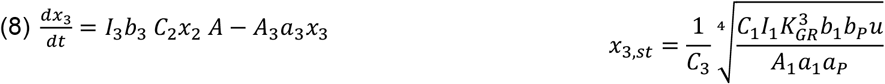

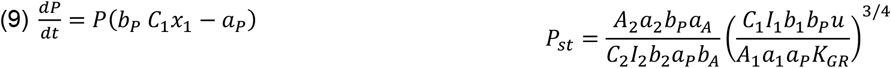

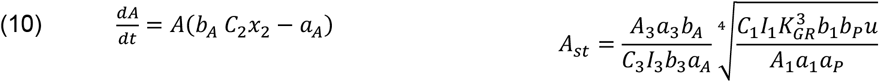

Under the approximation of *x*_3_ << *K*_*GR*_, the GR is not activated (*GR*(*x*) ≃ 1) and the steady state solution is:

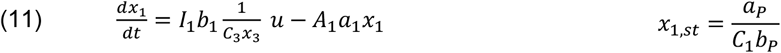

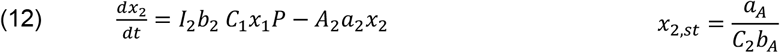

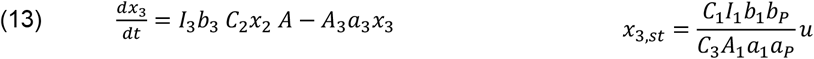

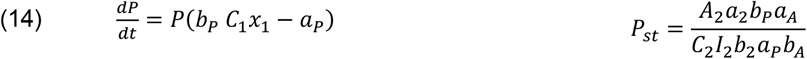

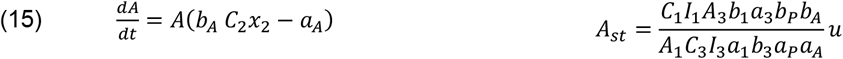

In both cortisol regimes, its steady state, *x*_3,*st*_, can be reduced only by CRH-synthesis inhibitors, *I*_1_ < 1; anti-CRH-blocking antibodies, *A*_1_ > 1; and CRH-receptor antagonists, *C*_1_ < 1. Note that GR antagonists, *C*_3_ < 1, are predicted to increase cortisol levels in this scenario. However, the net effect of cortisol on target cells, *C*_3_*x*_3_, after accounting for the competing antagonists (*C*_3_), would cancel out. The steady state solution also shows that CRH-receptor antagonists (*C*_1_ < 1) are predicted to increase CRH levels, and thus less favorable.

These results do not hold in a non-adjustable gland mass model (see Supplementary information).

#### Pituitary adenoma (Cushing’s Disease)

To model a pituitary adenoma, we use the approach developed by ^64^. We add the tumor ACTH secretion capacity, *T*_*P*_(*t*), to equation (2). We assume the tumor increases with time according to a logistic rule. The exact form of tumor growth does not influence our conclusions because, as demonstrated below, the bifurcation of the system’s steady state depends on the tumor crossing a specific threshold and not on its exact temporal dynamics.

To study the disease progression we start with normal levels of cortisol which on average obey*x*_3_ << *K*_*GR*_. In this limit *GR*(*x*) ≃ 1 and the system becomes:

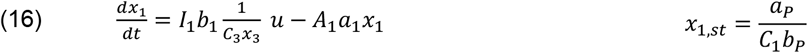

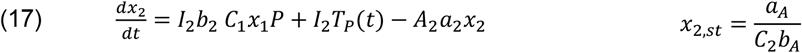

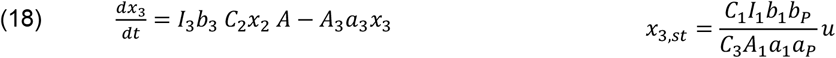

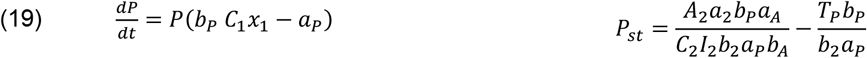

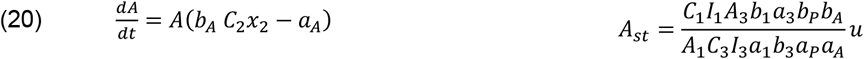

As long as the tumor secretion is below a threshold, 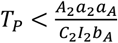, the steady states of all HPA components stay fixed except for the pituitary, which compensates and decreases with tumor size. This regime corresponds to the subclinical phase of Cushing’s disease. When the tumor is big enough to cross the threshold 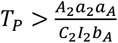, the pituitary mass goes to zero. In this limit, *P* → 0, ACTH steady state depends only on tumor secretion, 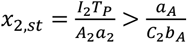 (from equation (17)). Substituting this lower limit of *x*_2,*st*_ in equation (20) for adrenal growth, we obtain a positive net growth rate which means uncontrollable growth of the adrenal.

Limiting adrenal growth by a carrying capacity, *K*_*A*_, stabilizes the system. Here is the steady state solution of the pituitary adenoma system with an adrenal carrying capacity in the limit where *P* → 0.In this regime, cortisol levels are high, and thus we assume *x*_3_ >> *K*_*GR*_:

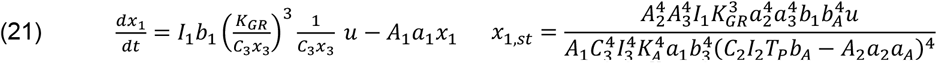

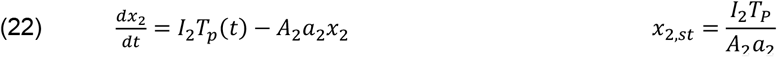

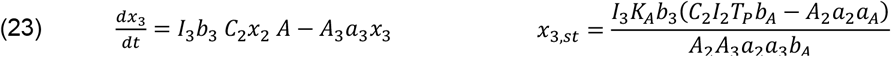

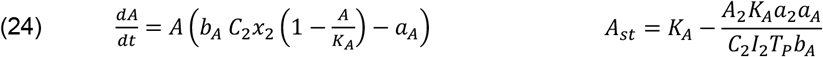

We learn from *x*_3,*st*_ equation that interventions targeting ACTH or cortisol (*I*_*i*_, *A*_*i*_, *C*_*i*_: *i* = 1,2) alter cortisol steady state whereas CRH-targeting (*i* = 1) interventions do not.

Note that as long as the pituitary is not completely suppressed (*P* > 0), ACTH-targeting interventions would increase the subclinical-clinical threshold 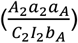. This will cause the system dynamics to flow back towards the subclinical regime. This transition also happens with cortisol-targeting drugs when adding adrenal carrying capacity in the first model (before assuming *P* → 0, equations (16)-(20)). However, the steady state of that model is not analytically solvable. We show this behavior with numerical simulations (Supplemental Figure 2).

#### Adrenal adenoma

We treat adrenal adenoma similarly to the way pituitary adenoma is modeled ^64^. We add the tumor cortisol secretion capacity, *T*_*A*_(*t*), to equation (3):

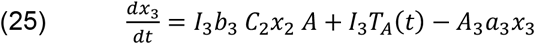

In this case the system has two steady states:

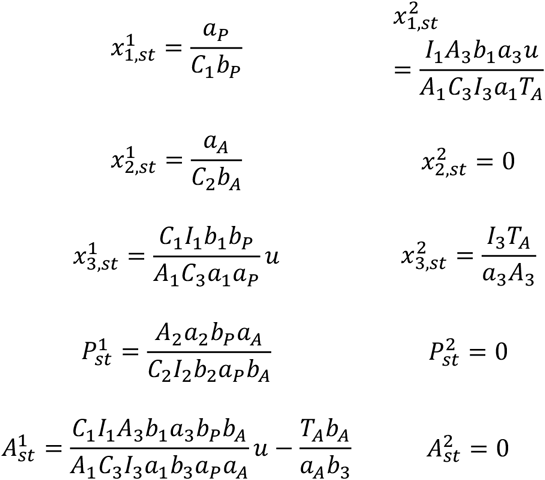

As long as the adrenal tumor secretion is below a certain threshold, 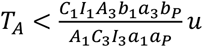, steady state is the only stable point of the system. This regime is the subclinical phase where all the HPA variables remain normal due to the healthy adrenal cortex that shrinks and compensates. When the tumor crosses that threshold, the adrenal cannot compensate further and it goes to mass zero, *A* → 0. Cortisol rises and CRH levels are too low so the net pituitary growth rate is negative and the pituitary mass goes to zero as well, *P* → 0. The system is drawn towards the second steady state, which becomes stable. Cortisol steady state is determined by tumor secretion capacity. Thus, in this case of adrenal adenomas, only cortisol-targeting drugs are predicted to work.

Cortisol-targeting drugs increase the subclinical-clinical threshold. If the glands are still able to recover, the system dynamics flow back to the subclinical regime. Note that CRH-modulating interventions could alter the threshold as well, however their effect will lead to an unwanted increase in long-term cortisol levels.

### Numerical simulations

To complete the system’s steady state solutions with analysis of its full dynamics, we used numerical simulations. Each simulation was initialized with the nominal HPA parameters ^21^ (Table 3) and a long warmup to reach the system’s steady state. This state, at the end of the warmup, was defined as the system healthy baseline before perturbation simulations. Then we ran a specific perturbation for each pathological condition: increased stressor input for the chronic stress conditions and growing tumor for Cushing’s syndrome. We simulated drugs as step functions administered during the clinical stage with a dose that predicts to cancel out the pathology effect, where possible. The results presented along this study are fold-changes of the HPA variables with respect to the healthy baseline.

**Table 3.** Parameters of the HPA gland-mass model.

| Parameter | Value | Reference |
| --- | --- | --- |
| $a_1$ | 0.17 [1/min] | 60 |
| $a_2$ | 0.035 [1/min] | 60 |
| $a_3$ | 0.0086 [1/min] | 60 |
| $a_P$ | 0.1 [1/day] | 21 |
| $a_A$ | 0.05 [1/day] | 21 |
| $K_{GR}$ | 4 | 21 |
| $n$ | 3 | 21 |

Numerical simulations were implemented in python 3.9.7 using solvers of ordinary differential equations (ODEs), implemented in the scipy package version 1.7.3. ^73^.

## Code and data Availability

Python code needed to reconstruct the analysis and figures is provided in the GitHub repository: https://github.com/tomermilo/hpa-drugs.

## Supplemental information

### HPA-targeting drug efficacy in non-adjustable gland model

Here we consider a model with glands that cannot change their functional mass. Thus, we set *P* = *A* = 1 and the system becomes:

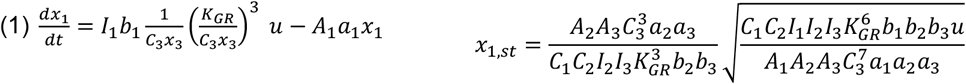

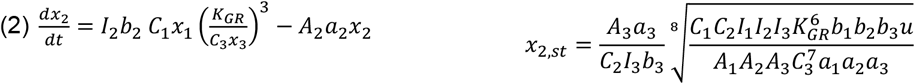

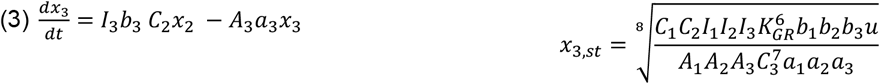

One can immediately see that according to this model all points of interventions should work to alter cortisol steady state. A complementary numerical simulation is depicted in Supplemental Figure 1.

### Full HPA dynamics under HPA-targeting drugs in Cushing syndrome

**Figure S1.**
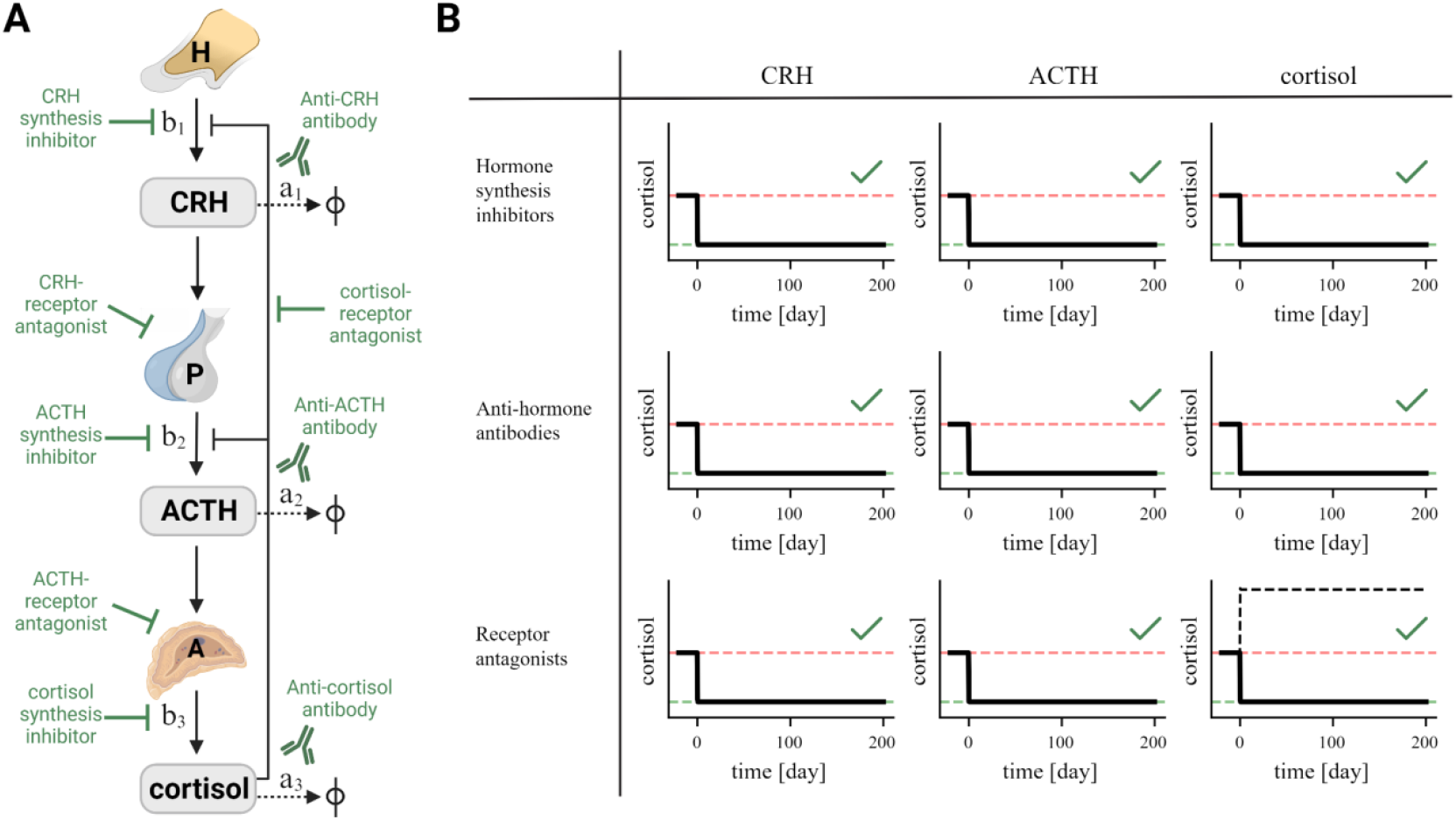
Efficacy of potential interventions according to a HPA model with non-adjustable glands, related to Figure 1. Panels similar to Figure 1.

**Figure S2.**
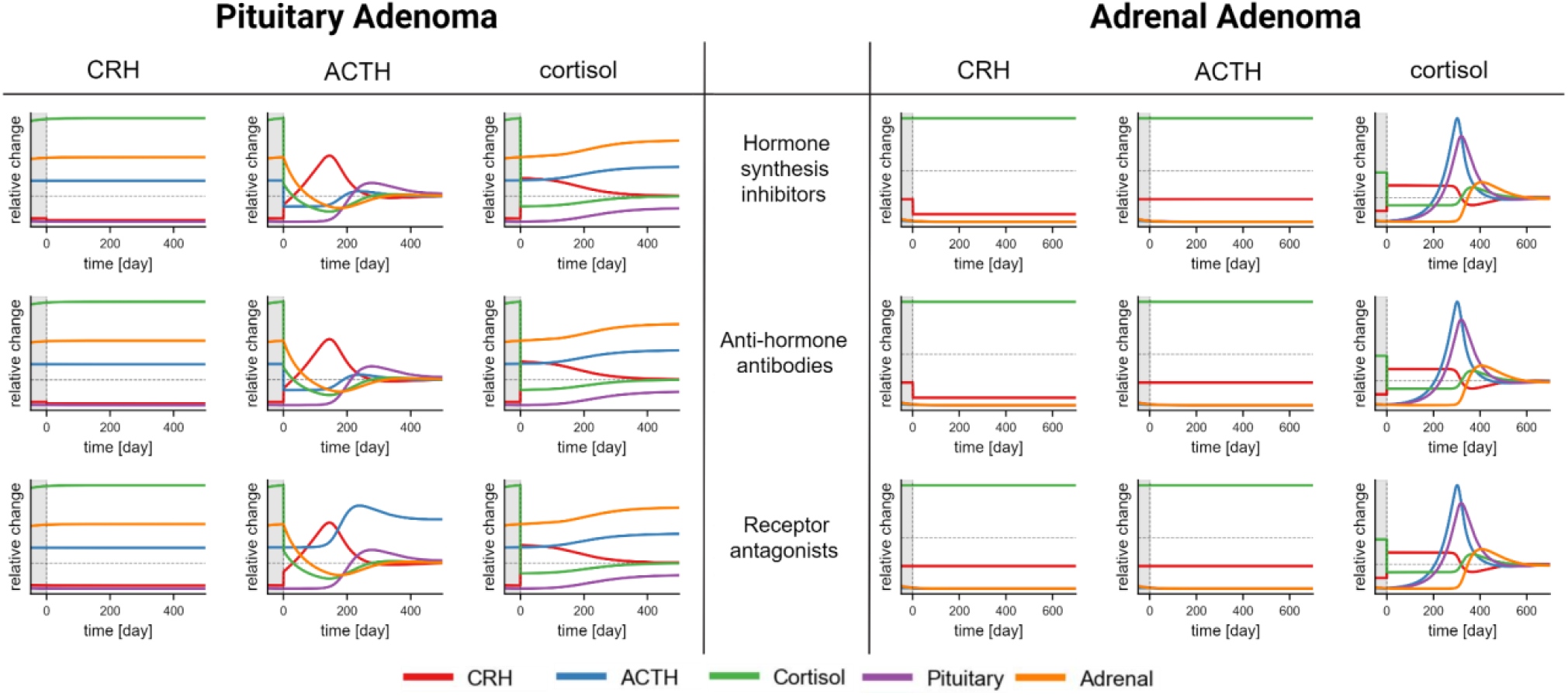
HPA dynamics under HPA-targeting drugs in Cushing syndrome, related to Figure 4. Simulations of the HPA dynamics during treating Cushing syndrome caused by a pituitary adenoma (left) or by an adrenal adenoma (right). The simulation starts with untreated Cushing (gray shaded region) and in time point zero the simulated drug is administered.

